# Towards Intelligent Intra-cortical BMI (i^2^BMI): Low-power Neuromorphic Decoders that outperform Kalman Filters

**DOI:** 10.1101/772988

**Authors:** Shoeb Shaikh, Rosa So, Tafadzwa Sibindi, Camilo Libedinsky, Arindam Basu

## Abstract

Fully implantable wireless intra-cortical Brain Machine Interfaces (iBMI) is one of the most promising next frontiers in the nascent field of neurotechnology. However, scaling the number of channels in such systems by another 10*X* is difficult due to power and bandwidth requirements of the wireless transmitter. One promising solution for that is to include more processing, up to the decoder, in the implant so that transmission data rate is reduced drastically. Earlier work on neuromorphic decoders only showed classification of discrete states. We present results for continuous state decoding using a low power neuromorphic decoder chip termed <u>S</u>pike-input <u>E</u>xtreme <u>L</u>earning <u>Ma</u>chine (SELMA). We compared SELMA against state of the art <u>S</u>teady <u>S</u>tate <u>K</u>alman <u>F</u>ilter (SSKF) across two different datasets involving a total of 4 non-human primates (NHPs). Results show at least a 10% or more increase in the fraction of variance accounted for by SELMA over SSKF across the datasets. Furthermore, estimated energy consumption comparison shows SELMA consuming ≈ 9 *nJ/update* against SSKF’s ≈ 7.4 *nJ/update* for an iBMI with a 10 degree of freedom control. Thus, SELMA yields better performance against SSKF with a marginal increase in energy consumption paving the way for reducing transmission data rates in future scaled BMI systems.

## I. Introduction

Approximately 6 million in the US and roughly 1 in 50 people worldwide suffer from paralysis [1]. This amounts to a significant number of people dependent on assistance from caregivers for daily living. The need to restore function in these people to perform activities for daily living has motivated development of a host of assistive technologies. One of the most promising technologies is the intra-cortical Brain Machine Interface (iBMI). This technology involves sensing intra-cortical signals from the surface of the motor cortex which are then passed through several signal processing and decoding algorithmic blocks to drive effectors such as a computer cursor for communication [2], [3], robotic/natural arm for feeding [4], [5], wheelchair for locomotion [6] among others.

Despite these convincing demonstrations, significant problems persist in the adoption of these technologies for daily use. Firstly, majority of the present iBMI systems involve the use of transcutaneous wires to connect the implanted electrodes to external recording equipment. This leaves an opening in the skull thereby increasing the risk of infection. Secondly, the larger form factor and subsequent visibility of the current iBMI systems in the form of a pedestal on a patient’s head run the risk of raising cosmesis concerns among end users [7]. To counter both these problems, fully implantable wireless recording systems capable of transmitting raw neural data [8] have been reported as a viable solution. However, these systems require the use of large batteries making it challenging to implant [9]. Another major concern pertaining to these systems is the limited battery lifetime amounting to a few hours. For e.g., [8] reports battery lifetime of up to 7 hours, necessitating frequent recharging with concomitant overheating issues. This problem only worsens further with increasing channel count [10], [11] hindering scalability due to overheating [12] as well as hitting bandwidth constraints [13].

An alternative proposed architecture to address this issue [14] is to do signal processing up to decoding in the implant itself as indicated in Fig. 1(b). The underlying idea is that this approach reduces the transmitted data rate by orders of magnitude [14] leading to an increase in battery lifetime. A key point to note is that no such in-vivo hardware system has been reported so far. The work we present here is a key step towards building such a fully implantable system.

**Fig. 1.**
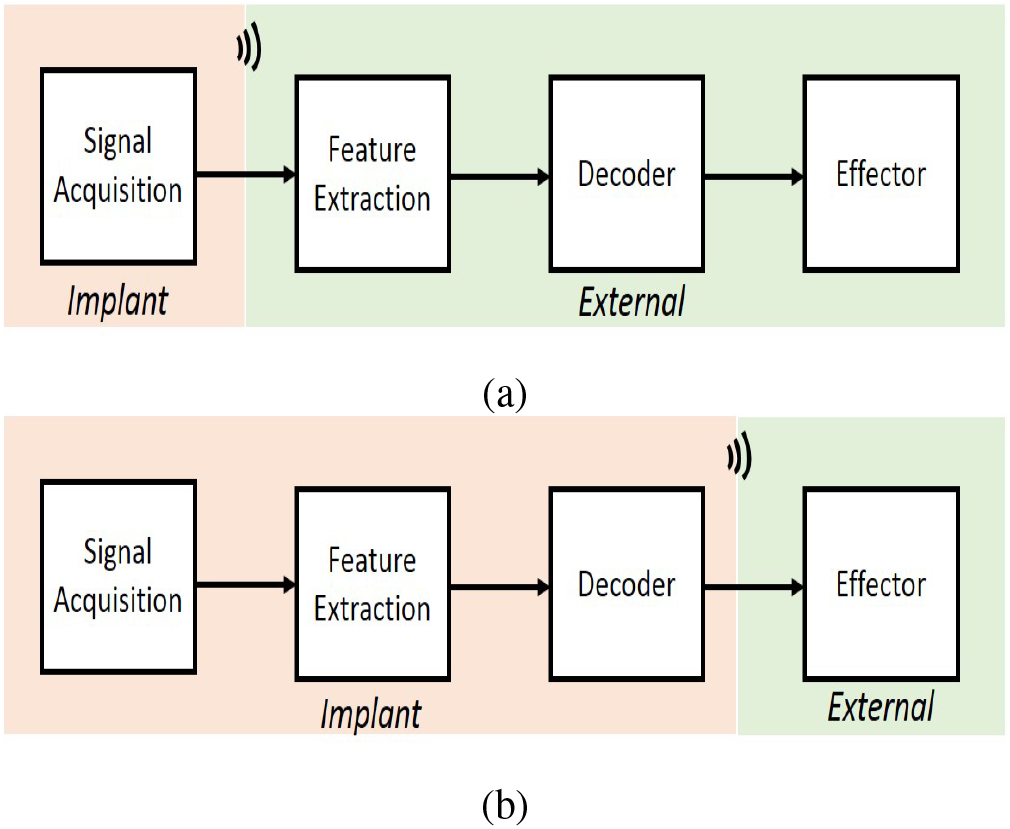
Block level representation of (a) conventional and (b) proposed wireless iBMI systems

In order to build such a system, signal acquisition, feature extraction and decoder blocks ought to be implemented in the fully implantable chip. Feature extraction block typically involves spike detection and low power implementations such as [15] consuming 40 *nW* /channel have been reported. While including a spike detector reduces bandwidth by 100*X* [14], the data rate still increases with increasing number of electrodes. The data rate can be made independent of the number of electrodes by integrating the intention decoder as well. However, the challenge in that case is to have sufficiently good decoding performance in limited power and area budget. [16] reports 6030 floating point operations per update for a Kalman Filter decoder consuming ≈ 1.82 *mW* on a ×86 processor for a 96 electrode array iBMI system driving a simple 2D cursor. The number of floating point operations are bound to dramatically rise with the increase in electrode densities and more degrees of freedom of control, ruling out direct mapping of these operations intensive algorithms on a fully implantable chip [16]. Thus, alternative computationally inexpensive algorithmic approaches have been reported in [16]– [21]. The lowest reported power is obtained by a neuromorphic approach that uses mixed analog-digital processing for a carefully chosen algorithm that exploits transistor mismatch for random weights [21]. However, this work only showed usage of the decoder in discrete classification tasks.

Depending on the mode of output, iBMI systems can be split into continuous mode and discrete mode systems. A discrete mode system is the one where the decoder outputs a discrete variable among a finite number of choices, an example being key selection on a keyboard [22]. A continuous mode iBMI system on the other hand outputs a continuous variable such as cursor’s velocity on a computer screen [23]. The continuous mode system is the more popularly used one owing to it’s generality of use in applications ranging from cursor control [2], [24] to robotic arm control [4] and paralyzed limb control [5].

The contributions in this work can now be summarized as follows:

- Presenting measured results from the micropower decoder chip in [21] for a continuous state decoding using publicly available datasets from 4 primates.
- Benchmarking the results against a Kalman Filter decoder to show consistently better performance.

Our results pave the way for widespread usage of randomized neural network based micropower decoders to scale future iBMI systems by the next factor of 10.

## II. Materials and Methods

We have used two different datasets to present our analysis and we will briefly describe these datasets in this section. The basic idea is to train a decoder to predict movement velocities from binned neural data.

### A. Experiment 1 (NHPs A,B)

This dataset is sourced from [25], [26]. It involves NHPs A, B doing a random target acquisition task through a planar manipulandum controlling a cursor being displayed on screen as shown in Fig. 2. Neural data acquisition was carried through a 96-channel Utah electrode array implanted in primary motor cortex (M1) and premotor cortex (PMd) areas in NHP A and primary somatosensory cortex (S1) in NHP B. 164 and 52 neurons were isolated in NHP A, B respectively. Datasets for NHP A, B correspond to 21 and 51 minutes respectively.

**Fig. 2.**
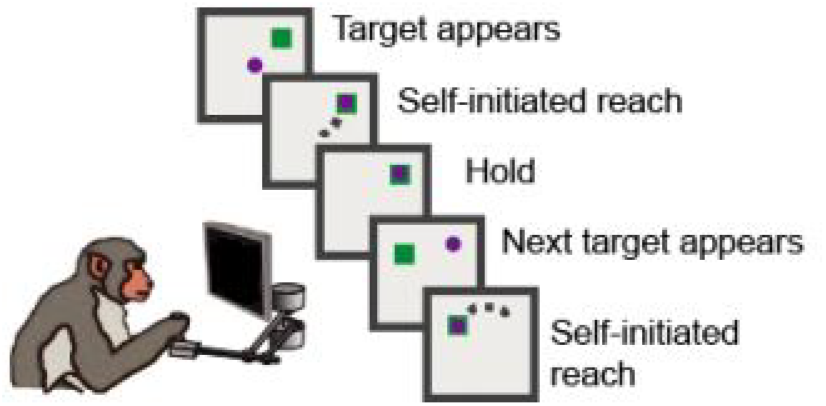
NHPs A, B control a cursor on screen through a planar manipulandum to reach targets flashed at random locations on a screen in a continuous manner one after another. A brief hold period is maintained between reaches. Image is obtained courtesy [25] (CC-BY license).

### B. Experiment 2 (NHPs C,D)

Data has been obtained courtesy [27]. NHPs C, D are trained to do self-paced reaching tasks without any time intervals. A virtual reality environment has been setup wherein the fingertip of NHP drives the cursor [28]. For more details of the experiment one is encouraged to read [28]. 96 channel Utah electrode array implanted in M1 area has been used for signal acquisition while an electromagnetic sensor is employed for tracking fingertip position. Recording for NHPs C, D lasts for up to 10 and 19 minutes respectively.

## III. Decoding methodology and algorithms

### A. Decoding paradigm

As mentioned earlier, we consider a continuous state decoding in this work. The input to an iBMI system at a given instance of time, *t* = *i* consists of an input feature vector 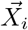 given as,

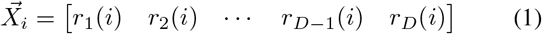

where 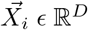 and *r*_*z*_(*i*) is computed as the number of spikes occurring at the *z*^*th*^ electrode looking back in a time bin - *T*_*bin*_ ms. For every 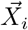, the continuous mode decoder outputs a continuous variable 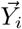, where 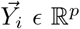 and *p* represents the degrees of freedom of control. Different algorithms employ different techniques to learn the mapping from 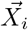 to 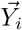. State of the art variable 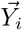 employed in iBMIs today is the velocity of the effector [2], [16], [24], [30], [31] and hence the decoder is often referred to as velocity Kalman filter (vKF). We will first briefly describe the state of the art technique - Kalman filter followed by a neuromorphic alternative.

### B. Algorithms

#### 1) State of the art - Kalman Filter

Kalman filter [32], [33] and it’s variants such as unscented Kalman filter [34], [35], steady state Kalman filter [36] have been the workhorse of decoder algorithms in iBMI systems. The proponents of this technique argue for its use over other techniques such as population vector [37], linear filtering [38] algorithms based on its sound probabilistic foundation, and the fact that it utilizes information over time in a recursive fashion [33]. Kalman filter models linear relationship between 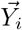 (kinematic state) and 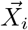 (neural observations) at time *t* = *i* as follows:

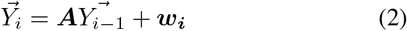

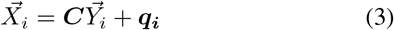

where ***A*** ϵ ℝ^*p×p*^ represents the state matrix, ***C*** ϵ ℝ^*D×p*^ represents the observation matrix, ***w***_***i***_ ϵ ℝ^*p×p*^ and ***q***_***i***_ ϵ ℝ*D×D* are Gaussian noise sources and are defined as ***w***_***i***_ ~ 𝒩(0, ***W***) and ***q***_***i***_ ~ 𝒩(0, ***Q***). Decoder training involves learning the parameters ***A**, **C**, **W**, **Q*** from training data.

At time step *t* = *i*, an a priori kinematic state estimate 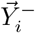 from previous time step *t* = *i −* 1 is computed as,

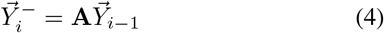

followed by computation of error covariance matrix 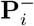, where 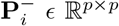,

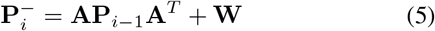

Using the measured firing rate 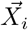 and the estimated a priori kinematic state 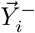, the final estimated kinematic state 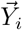 is obtained as,

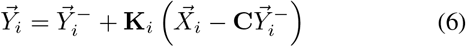

followed by computation of posterior error covariance matrix **P**_*i*_ ϵ ℝ^*p×p*^,

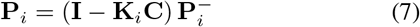

where **K**_*i*_ ϵ ℝ^*p×D*^ is the Kalman gain matrix and is computed as,

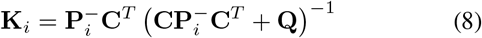

A simpler lower complexity alternative - the steady state Kalman filter (SSKF) has been reported in [36]. In terms of performance, there is almost no deficit as the correlation coefficient between decoded velocities from steady state and original Kalman filter has been reported to be as high as 0.99 [36]. Kalman gain is not updated at every time instance and instead held fixed as,

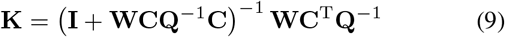

The kinematic state 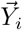 is updated based on the below equation,

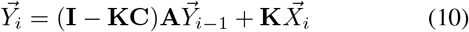

#### 2) A Neuromorphic Alternative - Extreme Learning Machine

A neuromorphic electronic system is one in which elementary physical phenomena in the form of analog signals are used as computational primitives rather than discrete digital signals [39]. The benefit of this approach is that it can lead to orders of magnitude of reduction in power consumption for pattern recognition applications [39], [40]. As far as neuromorphic compatible learning algorithms are concerned, the neural engineering framework (NEF) [41], [42] is a popular choice. The NEF essentially encodes the inputs in a non-linear fashion using random projections, which are then subsequently linearly decoded in order to model an arbitrary function. [16], [43] reports comparison of velocity Kalman filter (vKF) against NEF in a center-out iBMI task involving NHPs. A related algorithm, developed independently in the machine learning community, is the extreme learning machine which is akin to NEF without the recurrent connections. Briefly put, ELM is a single hidden layer feedforward neural network involving fixed, random non-linear projections in the first layer followed by a single-shot learning of weights in the second layer [44], [45]. For an input feature vector 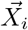 defined in Equation (1), the corresponding hidden layer vector is given as,

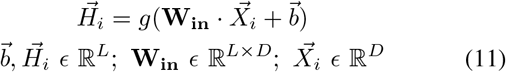

where *g*(·) stands for the activation function, **W**_**in**_ represents the input layer weights, *L* denotes the number of hidden neurons and 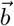 denotes the bias values associated with hidden layer neurons.

Output 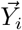 corresponding to *X*_*i*_, *H*_*i*_ is given as,

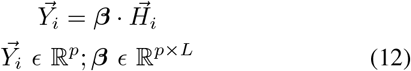

where ***β*** represents the second layer weights and *p* is number of degrees of freedom of output control.

Since the first layer weights are fixed, the optimal output weights ***β*** can be computed in a single step as shown below,

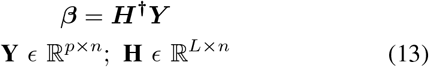

where ***H***^**†**^ represents the Moore-Penrose inverse of **H**. Thus, single step calculation of output weight lends to extremely fast learning while skirting around problems of local minima typically faced in gradient-based learning algorithms like backpropagation [45].

#### 3) SELMA - A Neuromorphic Processor

SELMA [21] is an ultra-low power hardware implementation of ELM with 128 channel input capacity as shown in Fig. 3(b). It takes inputs in the form of spikes and has the provision of configuring bin-width to compute firing rates. It exploits inherent device mismatch in a current mirror to provide fixed first layer random weights, thereby exploiting it as a single transistor multiplier. Second layer multiplication has been implemented on a DSP in the initial version [21] and subsequent versions [29], [46] have reported an integrated second stage on the same chip. For more details, interested readers are encouraged to refer to [21], [29], [46].

**Fig. 3.**
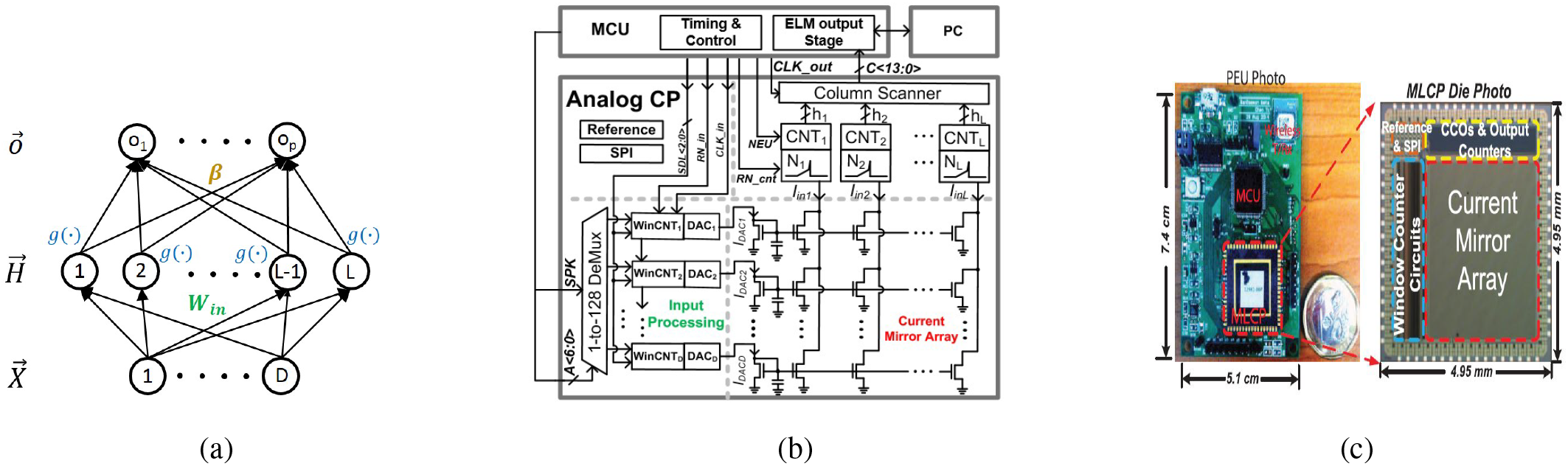
(a) ELM neural network architecture with 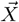, 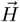 and 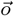 representing input, hidden and output layers respectively. **W**_**in**_, ***β*** represents first and second layer weights while *g*(·) stands for the activation function, (b) ELM chip-level architecture [21] is presented here. Firing rates are computed from spikes received as inputs by input window counters. Digital-Analog converters translate the firing rates into currents in each input channel. Current mirror array is used to realize first layer random projection. Current controlled oscillator is employed at the hidden nodes to yield a value proportional to the current at the respective hidden node. Second layer multiplication is implemented on a MSP430 microcontroller (MCU) in this work but can be integrated on-chip as well [29], (c) Picture of the test board in the form of a Portable External Unit (PEU) made up of machine learning co-processor (MLCP) and MCU is presented here [21]. MLCP implements the first layer of computation while MCU is employed for second layer computation.

## IV. Measured Results

### A. Analysis Methodology

We present comparison of KF, SSKF and SELMA across the experiments reported in section II as they produce decoded 2D components of movement velocities at 10 Hz. Bin-width for computation of firing rate is typically chosen as few hundreds of ms. Accordingly, we have compared the methods while sweeping bin-widths in steps of 100 ms. Decoder performance is also known to be subject to the amount of training data used to train a model. Thus, models were trained on varying amounts of training data while reporting performance on held-out test data. Electrodes with median firing rates less than 1 Hz were excluded from analysis.

The fraction of variance accounted for (FVAF) is used as a metric of comparison over Pearson’s correlation coefficient following the methodology used in [26], [47]. [47] reports it to be a superior and stricter goodness of fit indicator than correlation coefficient as it requires an ideal match between predicted and true values than simple correlation. FVAF is formulated as:

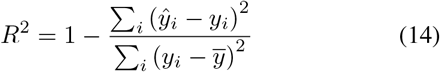

where 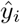, *y*_*i*_ and 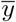 stand for predicted, true and mean values of the output variable. *R*^2^ is computed for each of the 2D components of movement velocity and its average value is plotted in Fig. 4 and Fig. 5.

**Fig. 4.**
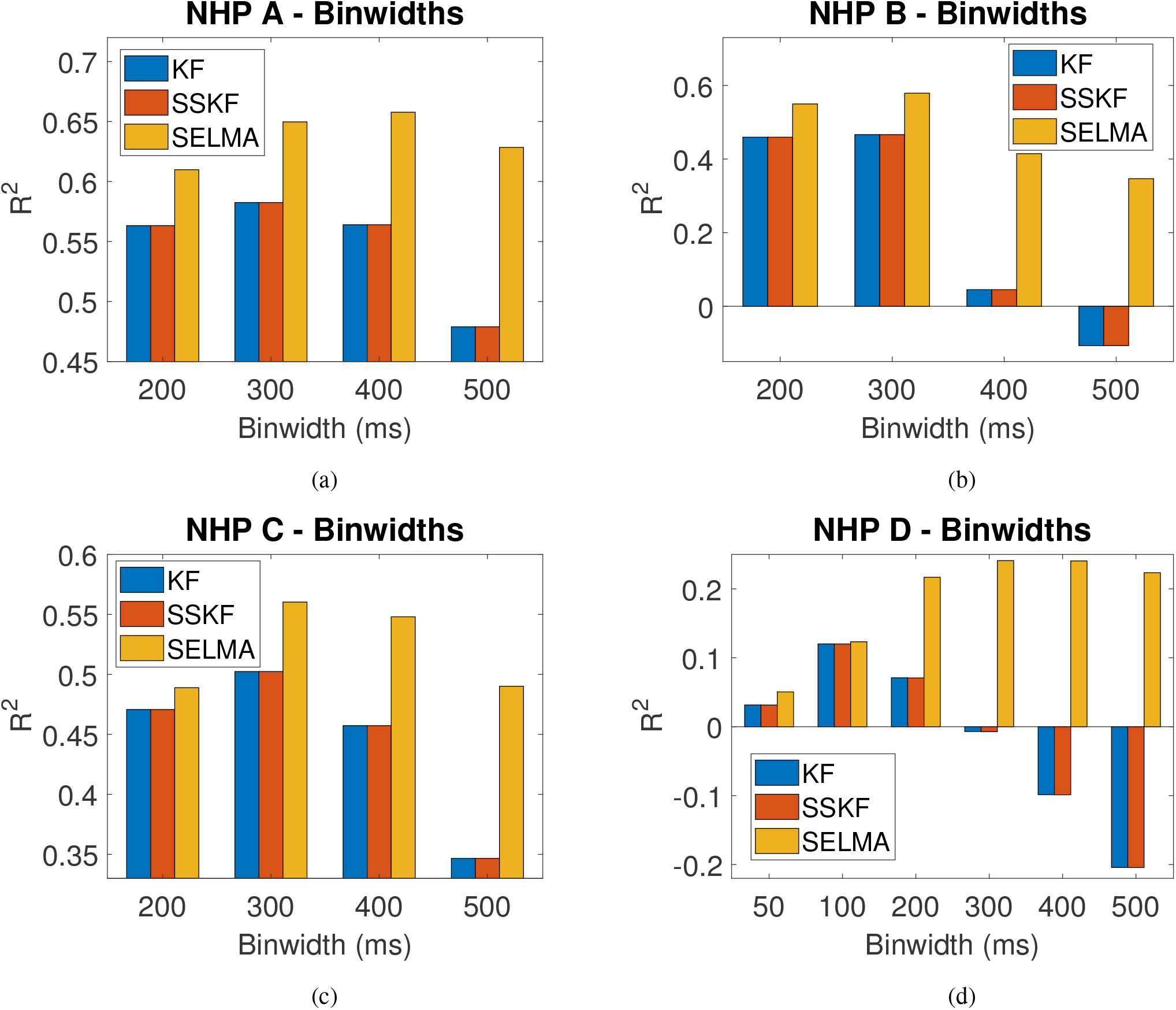
KF, SSKF and SELMA models are trained on the first half period of respective datasets and the remaining half is used as testing set. Plots show decoding results in the form of averaged *R*^2^ across 2D components of movement velocity, as bin-widths are swept for computing firing rates. The proposed neuromorphic decoder SELMA exhibits comparable or higher *R*^2^ across all bin-widths and datasets proving the robustness of our method.

**Fig. 5.**
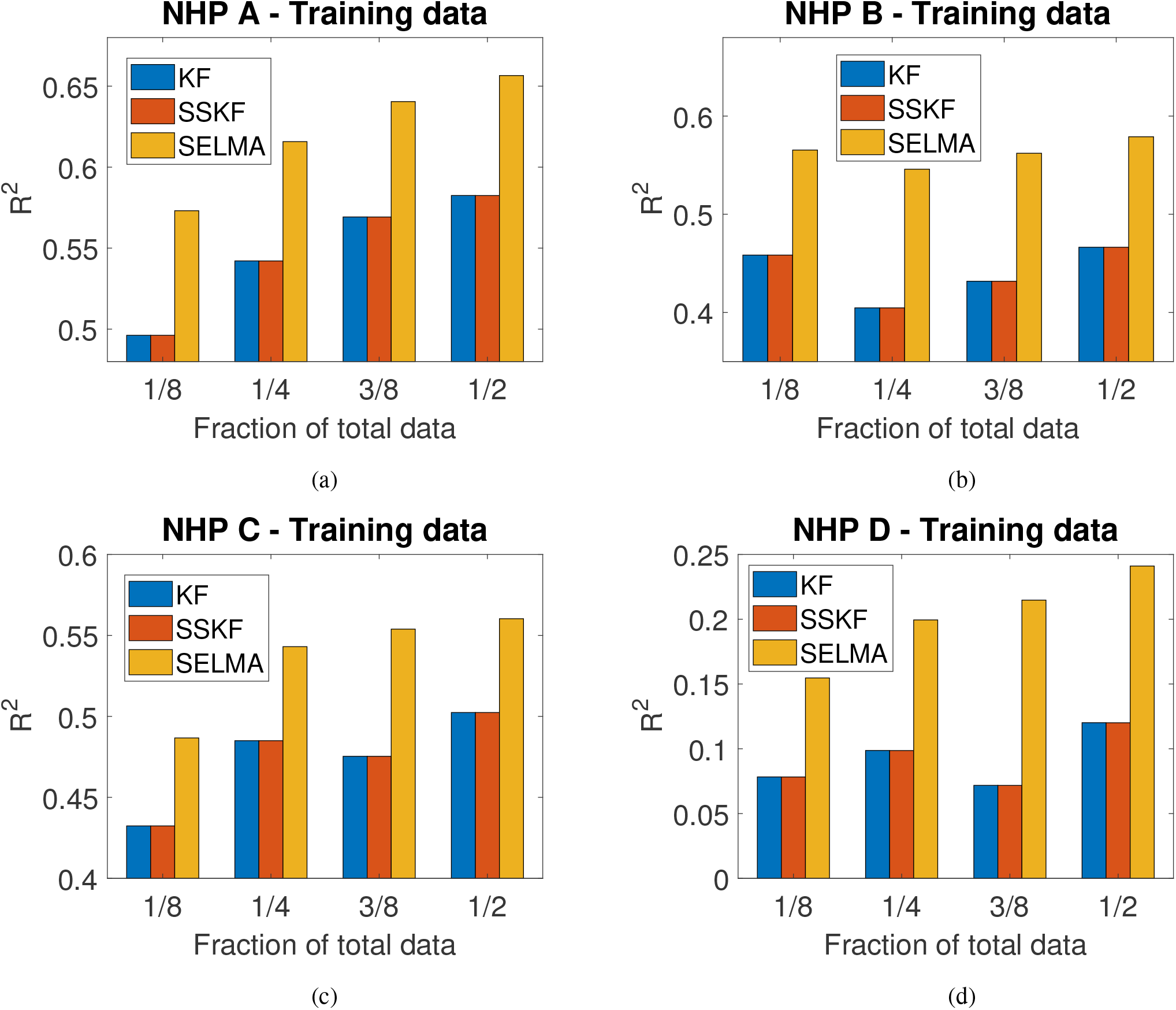
Plots show impact of amount of training data on decoding capability. For each decoder, bin-width corresponding to highest *R*^2^ in Fig. 4 is used for computation of firing rates. The proposed neuromorphic decoder outperforms both KF and SSKF decoders on all four datasets.

### B. Experimental Setup

The experimental setup in this work involved passing pre-recorded spikes through a custom MATLAB program written on a PC to the input channels of the chip and reading back hidden layers on the PC. During the training phase, read back hidden layers (**H**), target matrix (**Y**) were employed to obtain second layer output weight (***β***) following Equation. 13. A key difference to note is that SELMA was employed to build a regression model in this work as compared to a classification model reported in the earlier works [21], [48], [49]. During the testing phase, read back hidden layers (**H**) were multiplied by ***β*** as per Equation. 12 to arrive at the decoded output 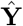. Although we are presenting offline results here, SELMA has been demonstrated to operate in a real-time closed loop setup involving a discrete-mode iBMI [49] and can easily be extended to continuous-mode iBMI as well.

### C. Results

#### 1) Sweeping bin-widths

Fig. 4 shows comparison of KF, SSKF and SELMA as the respective decoders are trained on the first half of NHP A, B, C and D datasets and tested on the remaining half. The performance of KF and SSKF is almost identical along the lines of previously reported results [36]. SELMA outperforms KF, SSKF by accounting for *≈* 14%, 24%, 12%, 100% more variance than that obtained by KF, SSKF for NHPs A, B, C, D respectively.

#### 2) Sweeping training times

Fig. 5 shows comparison of KF, SSKF and SELMA as the respective decoders are trained on varying amounts of training data across datasets with the optimal bin-width for each decoder obtained in Fig. 4. One can see SELMA consistently outperforming KF, SSKF across NHPs and varying amounts of training times in Fig. 5. Maximum improvements of ≈ 15%, 35%, 16%, 200% across NHPs A, B, C and D respectively. Furthermore, one can observe decoding accuracy declining with increase in the amount of training data in some cases and is prominently observed in NHP B for KF, SSKF. This counter-intuitive result is due the non-stationary nature of neural data [50]– [54]. However, SELMA appears to be relatively robust against non-stationarity along the lines of similar results obtained previously in a discrete control paradigm in [48]. Furthermore, a visual comparison of decoding capability of different algorithms is presented in Fig. 6.

**Fig. 6.**
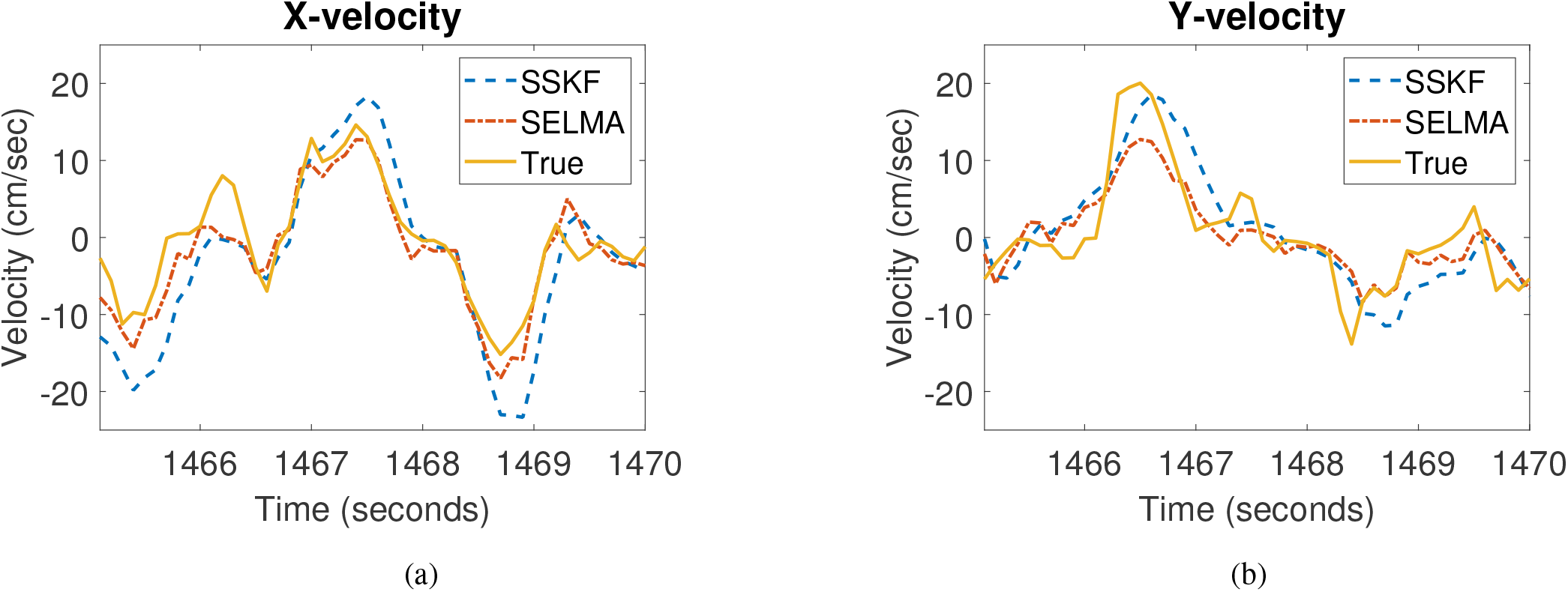
Example segment of decoded 2D components of velocities in NHP B comparing SSKF and SELMA decoders. SELMA outperforms SSKF in the above time snippet of *≈* 5 seconds.

### D. Comparison of Computational Complexity

In Fig. 4 and Fig. 5, we observe that KF and SSKF yield almost identical performance across NHPs and varying values of bin-widths and training data. Despite being more computationally complex, KF does not proffer any improvement over SSKF in terms of decoding capability. Thus, in Table. I, we present a computational complexity comparison of only SSKF against SELMA. The number of hidden neurons is a hyperparameter which is selected based on a validation set. As an example, Fig. 7 shows this by way of *k − fold* cross-validation for NHP A.

**Fig. 7.**
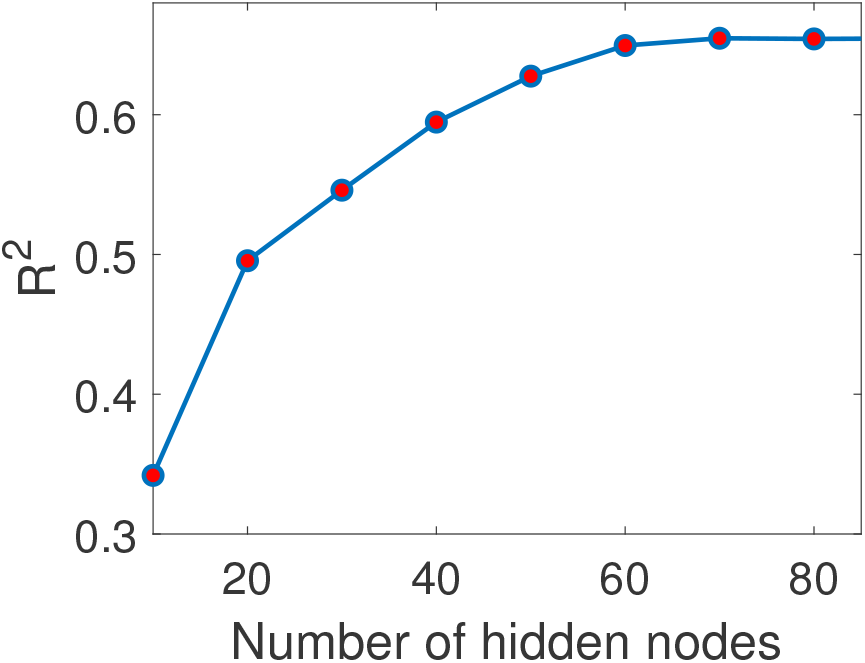
Validation accuracy is plotted for varying values of number of hidden nodes for NHP A’s training data with 64 input channels. *R*^2^ saturates around 70 hidden layer neurons in this case.

**TABLE I.** Computational Complexity Comparison for a 2-D iBMI decoder

| Algorithm/<br>Hardware | Analog<br>MACs | Digital<br>MACs | Energy/<br>update($nJ$) | Power<br>dissipation ($nW$) |
| --- | --- | --- | --- | --- |
| SSKF | - | 132 | 1.32 | 13.2 |
| SELMA | 4480 | 140 | 3.42 | 34.2 |

MAC = multiply and accumulate, Energy/(Analog MAC) = 0.45 pJ [21], Energy/(Digital MAC) = 10 pJ [55], Decoder update frequency = 10 Hz, Number of input channels, $D = 64$ , Number of hidden layer nodes in SELMA, $L = 70$ and Number of outputs, $p = 2$ .

In order to compare computational complexity, we consider the number of input channels as, *D* = 64 and the number of decoded outputs as, *p* = 2. For SELMA, we consider, number of hidden layer neurons as, *L* = 70 based on the value obtained in Fig. 7 for NHP A with 64 input channels.

For the decoder considered in Table. I and operated at 10 Hz, the power consumption comes to be around 13.2 and 34.2 *nW* for SSKF and SELMA respectively. Furthermore, if we consider a relatively advanced case of a decoder with a 10 degree of freedom output to control an anthromorphic arm [53] with the remaining decoder parameters same as Table. I, the energy consumption comes to be ≈ 9 *nJ/update* and 7.4 *nJ/update* for SELMA and SSKF respectively. Thus, one can conclude that SELMA consumes energy in a comparable range to SSKF. The reason for energy consumption of SELMA to be in the same realm as SSKF, despite overall greater number of operations, is the parsimonious analog implementation of the first layer. For e.g., if ELM were to be implemented using conventional digital hardware, energy consumption for the aforementioned case would come to be significantly greater around *≈* 46.2 *nJ/update*.

## V. Discussion

Kalman filter models linear relationship between input and output variables and works best when the input noise is Gaussian. However, in practice often the assumptions of linearity and Gaussian nature of recording noise don’t hold true in case of iBMIs [28]. ELM on the other hand, by virtue of random projection translates the input into a non-linear higher dimensional space, thereby increasing the generalization ability of the model as stated in Cover’s theorem [56].

In fact, there have been recent initial offline studies which have shown even more complex non-linear algorithms such as Long Short Term Memory (LSTM) neural networks out-performing SSKF [31], [57]. However, we can observe three critical shortcomings in an LSTM-based approach. Firstly, it requires a larger amount of training data. For e.g., [57] reports using data spanning around 73 calendar days while using LSTM. Secondly, complex neural networks are known to overfit to noise and optimally tuning the hyper-parameters is often a daunting impediment to overcome [58]. Third and most importantly, these neural networks are much more computationally complex than both SSKF and SELMA. This results in higher training times and often dedicated hardware such as graphic processing units (GPUs) for training, making the system cumbersome to use [59]. Furthermore, the amount of MACs involved for the LSTM network presented in [57] to control a 2D cursor comes to ≈ 22900. This is around two orders of magnitude more computationally complex than SSKF, SELMA and possibly rules out the prospect of implementing the decoder in the implant itself. Note that we have considered the same input and output conditions for LSTM as was used in Table. I.

Electrophysiological signal acquisition techniques are advancing expeditiously with the latest offering of Neuropixels probe [11] capable of recording up to 1000 neuron channels. [10] reports a Moore-like law with the number of simultaneous neurons recorded doubling every ≈ 7 years over the last 5 decades. We consider a case of an iBMI controlling a 10*D* anthromorphic arm at an operating frequency of *f*_op_ = 50 Hz. For such an application, we compare the resulting transmission data rates across systems transmitting a) raw data, b) firing rates and c) decoded outputs for varying number of input channels (*N*_chan_) in Fig. 8. Systems transmitting firing rates involve integration of a spike detector followed by a counter to yield firing rates at frequency - *f*_op_ for every input channel. In addition to the spike detector and counter blocks, systems transmitting decoded outputs involve integration of the decoder block as well in the implant as seen in Fig. 1(b). Transmission data rates for the different cases can be computed as per the formulation below,

**Fig. 8.**
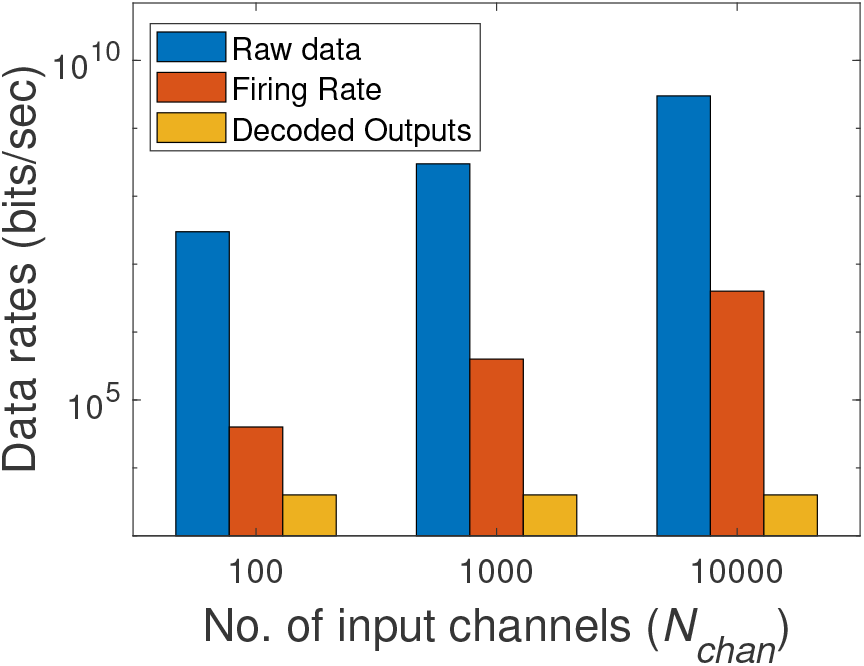
Comparison of transmission data rates between systems transmitting a) raw data, b) firing rates and c) decoded outputs for a 10D iBMI operating at 50 Hz. It can be seen that the system with decoded outputs is more scalable since its output data rate is orders of magnitude lower than both other options.

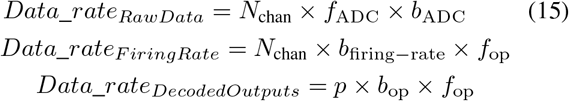

where, calculation considerations in Fig. 8 involve ADC Sampling rate (*f*_ADC_) = 30 kHz, ADC bit resolution (*b*_ADC_) = 10, Firing rate bit resolution (*b*_firing*−*rate_) = 8, iBMI operation frequency (*f*_op_) = 50 Hz, Number of outputs (*p*) = 10 and Decoded output bit resolution (*b*_op_) = 8.

Thus, one can observe that integrating signal processing blocks beyond conventional signal acquisition block in the implant leads to at least 3 or more orders of magnitude of transmission data rate reduction. The system transmitting decoded outputs is of particular significance since it makes the transmission data rate essentially agnostic to the number of input channels. However, one can be rightfully concerned about the increase in the power consumption due to - a) addition of firing rate computation block in system transmitting firing rates and b) addition of firing rate and decoder blocks in system transmitting decoded outputs. To gain an understanding about the amount of overall system level power saving, let us again consider a 10*D* iBMI operating at 50 Hz with *N*_chan_ = 1000. Data rate was computed using formulation in Equation. 15 and transmitter power dissipation was found by multiplying data rate to reported transmitter power efficiency of 3.775 nJ/bit [8]. In the firing rate computation block, only spike detection block was considered for simplicity with 40 nW/channel [15] power consumption. Finally, SELMA was employed as a decoder considering reasonably large number of hidden layer neurons, *L* = 1500, *Analog MACs* = 15 × 10^6^, *Digital MACs* = 15 × 10^3^, *Energy/*(*Analog MAC*) = 0.45 pJ [21], *Energy/*(*Digital MAC*) = 10 pJ [55]. The normalized power dissipation figure in the last column of Table. II shows significant amount of power savings attained by virtue of incorporating both firing rate extraction and decoder blocks in the implant compared to the other two cases, thereby increasing battery lifetime.

**TABLE II.** Adding signal processing in the implant

| Systems Output | Data rate (kbps) | Transmission power (W) | Added blocks power dissipation (μW) | Power dissipated normalized to Decoded Outputs system* |
| --- | --- | --- | --- | --- |
| Raw Data | $3 \times 10^5$ | 1.13 | - | 11728 |
| Firing rate | 400 | 1.5m | 40 | 16 |
| Decoded Outputs | 4 | 15.1μ | 81.25 | 1 |

\*Power dissipation of all the blocks post signal acquisition in the implant is considered, Wireless transmitter power dissipation = 3.775 nJ/bit [8], $N_{chan} = 1000$ , (Spike-detector Power consumption)/(input channel) = 40 nW. SELMA is used as a decoder with $D = 1000$ , $L = 1500$ , $p = 10$ . Accordingly, decoder power consumption equals 41.25 $\mu$ W.

## VI. Conclusion and Future Work

We have presented hardware results of a neuromorphic chip - SELMA across different experimental setups and NHPs. Consistent improvement over state of the art technique - KF and SSKF has been achieved over different operating parameters of bin-widths and training data corpus. Furthermore, we have also presented an energy consumption comparison for different use cases and found SELMA to consume power in the same range as simple linear SSKF. The reduction in power consumption while implementing ELM on SELMA is primarily attributed to the ingenious trick of using ultra low-power analog computation technique in the first layer.

We intend to do a thorough closed loop comparison of the presented methods in the near future. Furthermore, in recent studies introducing recurrent connections have been shown to be effective in iBMIs. Keeping this in mind, we aim to evaluate hardware friendly methods such as reservoir computing [60] to check for benefits. On the hardware front, we intend to design an end-to-end fully implantable wireless iBMI system based on the results obtained so far. We believe our results pave the way for obtaining the next order of magnitude increase in channel count for iBMI systems by integrating intelligence in the implant.

## Acknowledgment

The authors would like to thank [25] and [27] for making the data publicly available. This work was supported through grant RG87/16 by MOE, Singapore.

